# Endogenous IL-10 Contributes to Wound Healing and Regulates Tissue Repair

**DOI:** 10.1101/2022.03.15.484452

**Authors:** Walker D. Short, Meredith Rae, Thomas Lu, Benjamin Padon, Tanuj J. Prajapati, Fayiz Faruk, Oluyinka O. Olutoye, Ling Yu, Paul Bollyky, Sundeep G. Keswani, Swathi Balaji

## Abstract

**Background:** Interleukin-10 (IL-10) is essential in fetal regenerative wound healing and likewise promotes a regenerative phenotype in adult dermal wounds. However, the role of endogenous IL-10 in postnatal dermal wound healing is not well established. We sought to determine the role of IL-10 in murine full thickness, excisional wounds that are splinted to prevent contracture and mimic human patterns of wound closure.

**Methods:** Full thickness, excisional wounds were made in wildtype (WT) and IL-10^-/-^ mice on a C57BL/6J background (F/M, 8wks old). In a subset of wounds, contraction was prevented by splinting with silicone stents (stenting) and maintaining a moist wound microenvironment using a semi-occlusive dressing. Wounds were examined for re-epithelialization, granulation tissue deposition, and inflammatory cell infiltrate at day 7 and fibrosis and scarring at day 30 post-wounding.

**Results:** We observed no difference in wound healing rate between WT and IL-10^-/-^ mice in either the stented or unstented group. At day 7, unstented IL-10^-/-^ wounds had a larger granulation tissue area and more inflammatory infiltrate than their WT counterparts. However, we did observe more F4/80^+^ cell infiltrate in stented IL-10^-/-^ wounds at day 7. At day 30, stented wounds had increased scar area and epithelial thickness compared to unstented wounds.

**Conclusions:** These data suggest that endogenous IL-10 expression does not alter closure of full thickness excisional wounds when wound hydration and excessive contraction are controlled. However, the loss of IL-10 leads to increased inflammatory cell infiltration and scarring. These data suggest that previous reports of increased rates of healing in IL-10^-/-^ mice ought to be revisited considering recent advances in wound healing models. Moreover, these new findings suggest that IL-10 contributes to regulation of inflammation without compromising the healing response.

## Introduction

The obligatory scar formation that results from healing of dermal injuries in postnatal mammals likely arose as an evolutionary adaptation to allow rapid wound healing in a contaminated environment.[1] However, patients and surgeons in the modern era place great emphasis on the cosmetic outcome of wounds and surgical incisions. Thus, the market for anti-scarring therapies continues to grow rapidly and is projected to reach a value of almost $35 billion by 2023, and almost double that by 2030.[2, 3] Beyond the cosmetic outcomes of wound healing, tissue fibrosis is a common endpoint in the pathogenesis of multiple chronic diseases, for which the skin serves as a convenient model system.[1, 4] [5] Thus, the accurate study of dermal fibrosis and the development of therapeutic advances to attenuate it are imperative.

Interleukin-10 (IL-10) is a well-studied cytokine produced by multiple cells in the innate and adaptive immune system. It plays a major role in the regulation of inflammatory cell and cytokine responses that may affect cutaneous wound healing. The innate immune system is first to be activated following cutaneous injury; damage associated molecular patterns (DAMPs) and pathogen associated molecular patterns (PAMPs) trigger the activation of and histamine release from mast cells, leading to monocyte and neutrophil recruitment to the wound.[6–8] IL-10, in concert with IL-4, has been shown to inhibit mast cell development, with a greater quantity of mast cells able to be derived from the bone marrow of IL-10^-/-^ mice compared to WT.[9] Other effects include the regulation of macrophage polarization and function, as well as regulation of pro-fibrotic cytokines such as transforming growth factor-β (TGF-β).[9–12] Shortly following the discovery that the midgestational fetus heals dermal wounds regeneratively without scarring, IL-10 was found to be elevated in fetal skin. Studies by our team and others have shown an essential role for endogenous IL-10 in the regulation of the fetal scarless phenotype via effects on extracellular matrix (ECM) production and inflammatory responses.[13, 14] Our group has since demonstrated that IL-10 overexpression in postnatal dermal wounds promotes regenerative healing in a dose-dependent manner through several mechanisms via a JAK/STAT3 pathway.[15, 16] Namely, IL-10 overexpression induces a pericellular matrix consisting of high molecular weight hyaluronan (HMW-HA) via fibroblast-specific signaling. This is critical to the regenerative phenotype in addition to its well-established role in the regulation of inflammatory cell and cytokine responses.

The laboratory mouse remains by far the most commonly used animal model for interrogation of the molecular, cellular, and physiologic mechanisms governing IL-10’s role in wound healing. The IL-10 knockout (KO) mouse allows investigation of the impact of this cytokine on dermal wound healing, fibrosis, and scarring outcomes for development of novel therapeutics.

Given their central role in our understanding of physiology, it is of great importance that the dermal wound models recapitulate human wound healing as technically, ethically, and financially as possible. Some aspects of murine physiology are simply distinct from that of humans. The panniculus carnosus (PC) is a thin layer of striated muscle located deep to the dermal adipose layer in mouse and other rodent skin, which functions to provide thermoregulation in unwounded skin, as well as contraction to aid in rapid wound closure.[17, 18] While the PC is present in lower mammals, it is a vestigial remnant in humans, found in the platysma of the neck.[19] In contrast, human wounds, when allowed to heal by secondary intention, rely on granulation and re-epithelialization rather than contraction of the PC to bring wound edges together. Thus, the study of wound healing in mice must control for the action of the PC if this research is to act as a valid translational model for human wound healing.

To this end, the splinted wound model using silicone rings to prevent wound margin contracture was introduced to better recapitulate the repair mechanisms underlying human wound healing.[20] This relatively simple and reproducible surgical model can be performed aseptically, and the wounds are covered with a semi-occlusive dressing to provide a moist wound environment considered to be beneficial to wound healing.[20, 21] The wound bed can be easily accessed for application of topical agents and the study of healing progress by photographic and intravital imaging. Harvested wounds can be examined histologically for both the epithelial gap (the distance between the healing epithelial wound margins), granulation bed, and remodeled scar characteristics (presence of various cell populations, vascularity, matrix alterations). Thus, this surgical mouse model is an essential tool that can aid researchers in closely representing physiologic human wound healing.

Previously published work showed that full thickness excisional dermal wounds on IL-10^-/-^ mice healed faster than WT, with an expected increase in alpha-smooth muscle actin (α-SMA) content and fibrosis.[22] However, the method of wounding used in this study did not account for the effect of the panniculus carnosus on wound closure. While no differences in mechanical strength were found between IL-10^-/-^ and WT scars, this contradicts results from an incisional wounding model in which IL-10^-/-^ mice heal with compromised mechanical integrity.[23] Furthermore, we have recently demonstrated that enriching wounds with type 1 regulatory T-lymphocytes, which are defined by their elevated production of IL-10, results in accelerated wound closure.[24] These contradictions highlight how the choice of murine wound models may significantly impact outcomes.

In this work, we have compared two models of full thickness murine wounds. The first includes wounds allowed to heal without stenting while maintaining a moist wound environment. In the second, initial wound contraction is prevented for 7 days with a silicone stent under a moist wound environment. We have evaluated the differences between these two models when applied to wound healing in WT and IL-10^-/-^ mice. We demonstrated that stenting wounds allows for a more human-like healing by secondary intention and results in a markedly different healing phenotype than unstented wounding. Moreover, stented wounds reveal that knockout of IL-10 does not impact epithelial closure.

## Materials and Methods

### Animal Studies

C57BL/6J (WT) and B6.129P2-Il10^tm1Cgn^/J (IL-10^-/-^) mice were purchased from Jackson Laboratory (Bar Harbor, ME), bred, and maintained under pathogen-free conditions with access to food and water ad libitum in the Texas Children’s Hospital Feigin Center animal facility. Protocols for animal use were approved by the Institutional Animal Care and Use Committee at Baylor College of Medicine.

### Wound Healing Model

Mice were anesthetized with 2-3% isoflurane via inhalation. Dorsal skin was shaved and prepared by scrubbing alternately with betadine and isopropyl alcohol for 3 times. Two full-thickness excisional wounds were created on the dorsum of 8-week-old WT or IL-10^-/-^ male and female mice using a 6-mm punch biopsy (Miltex, Plainsboro, NJ). *Stented group*: circular silicone stents with inner diameter 8mm and outer diameter 16mm were placed around the wound with a skin adhesive and sutured with 6-0 polypropylene suture in a simple interrupted fashion and maintained for 7 days post-wounding to prevent immediate contraction and allow for healing by secondary intention. *Unstented group*: No stents were applied to animals in this treatment group. Wounds of both treatment groups were covered with a sterile adhesive dressing (Tegaderm; 3M, St. Paul, MD) to allow healing in a moist protected environment. Wounds were harvested at days 7 and 30 for analysis. 5-6 mice were included in each treatment group at each time point.

### Histology and Immunohistochemistry

Skin tissue was harvested, fixed in 10% neutral buffered formalin and paraffin embedded. 5-μm thick sections were cut and mounted onto slides. Slides were deparaffinized and rehydrated to PBS following standard protocol and immuno-histochemistry (IHC) staining was performed on a Dako Auto-stainer Link 48 (DakoLink version 4.1, edition 3.1.0.987; Agilent, Santa Clara, CA). Primary antibodies against α-smooth muscle actin (α-SMA) for myofibroblasts (ab5694; Abcam, Cambridge, MA), CD31 for endothelial cells (ab28364, Abcam, Cambridge, MA), CD45 for pan leucocytes (ab10558; Abcam, Cambridge, MA), and F4/80 for pan-macrophages (ab111101; Abcam, Cambridge, MA) were detected by EnVision+System-HRP (DAB) kits (Dako North America, Carpinteria, CA) and hematoxylin counter staining. Histology slides were imaged with Leica DM 2000® with Leica Application Suite X® version 3.0.4.16529. Percentage of positive cells per high powered field (40x) within the granulating wound bed were quantified.

### Wound analysis

Wound healing response at day 7 and scar tissue morphology and ECM architecture at day 30 post-wounding were determined by hematoxylin and eosin (H&E) staining and Masson’s Trichrome Staining respectively (Leica Biosystems, Buffalo Grove, IL). Epithelial gap, granulation tissue, and collagen staining were quantified using ImageJ (National Institutes of Health, Bethesda, MD). Epithelial gap was measured as the distance between the epithelial wound margins of the newly forming epidermis at day 7. Granulation tissue and scar areas were measured as the area beginning from one end of the original wound margin, down to the ipsilateral PC, across the wound cleft to the opposite original wound margin, and finally back to the initial wound margin. Collagen content per HPF in the dermis of the scars was measured using established methods with color-thresholding in ImageJ in which color segmentation is used to isolate only blue pixels, representing collagen fibers, thus allowing quantification of the amount of collagen within the selected area.[25] Epidermal topography was measured as traced length of the undulating basal layer of the epidermis per 200μm of wound bed.

### Statistical Analysis

Statistical analysis of the data was performed using analysis of variance (ANOVA), followed by Bonferroni post hoc tests, or student t-test when appropriate using Prism9 (GraphPad Software, San Diego, CA). The data are expressed as the mean ± standard deviation. A p value of <0.05 was considered statistically significant.

## Results

### Stenting prolongs epithelial gap closure

We first sought to demonstrate the gross effect of stenting on rate of epithelial gap closure. Full thickness wounds in WT mice were either allowed to heal without stenting or with a circular silicone stent sutured in place to prevent primary contraction **(Figure 1a)**. Grossly at day 7, unstented wounds appear to close more rapidly than stented wounds. Even partial stent compromise prior to day 7 results in altered gross and histologic characteristics of the wound **(Figure 1a, red boxes)**. Broken stents earlier in the healing timeline, such as between days 3 and 5 post-wounding (stented wounds in row 2 and 5) led to a drastic difference in phenotype from other stented wounds, with a morphology more typical of an unstented wound. On the other hand, breakage of a stent later in the experiment, such as between days 5 and 7 post-wounding (stented wound in row 3) led to a phenotype in between those with no stent and those that retained their stent for all 7 days post-wounding. Epithelial gap (the distance between neo-epidermal tongues) and granulation tissue area were quantified and compared between the unstented and stented groups that retained intact stents until day 7 post-wounding **(Figure 1b)**. Unstented WT wounds healed significantly more rapidly than wounds in which stents remained intact for a full 7 days post-wounding (unstented 1.16±0.71 mm vs Stented 4.13±1.72 mm, p<0.001). Stented wounds also had much smaller granulation tissue area when compared to unstented wounds at this time point (unstented 0.84±0.38 mm^2^ vs Stented 0.41±0.08 mm2, p<0.05). These data suggest that stenting prolongs wound healing time of murine dermal wounds.

**Figure 1:**
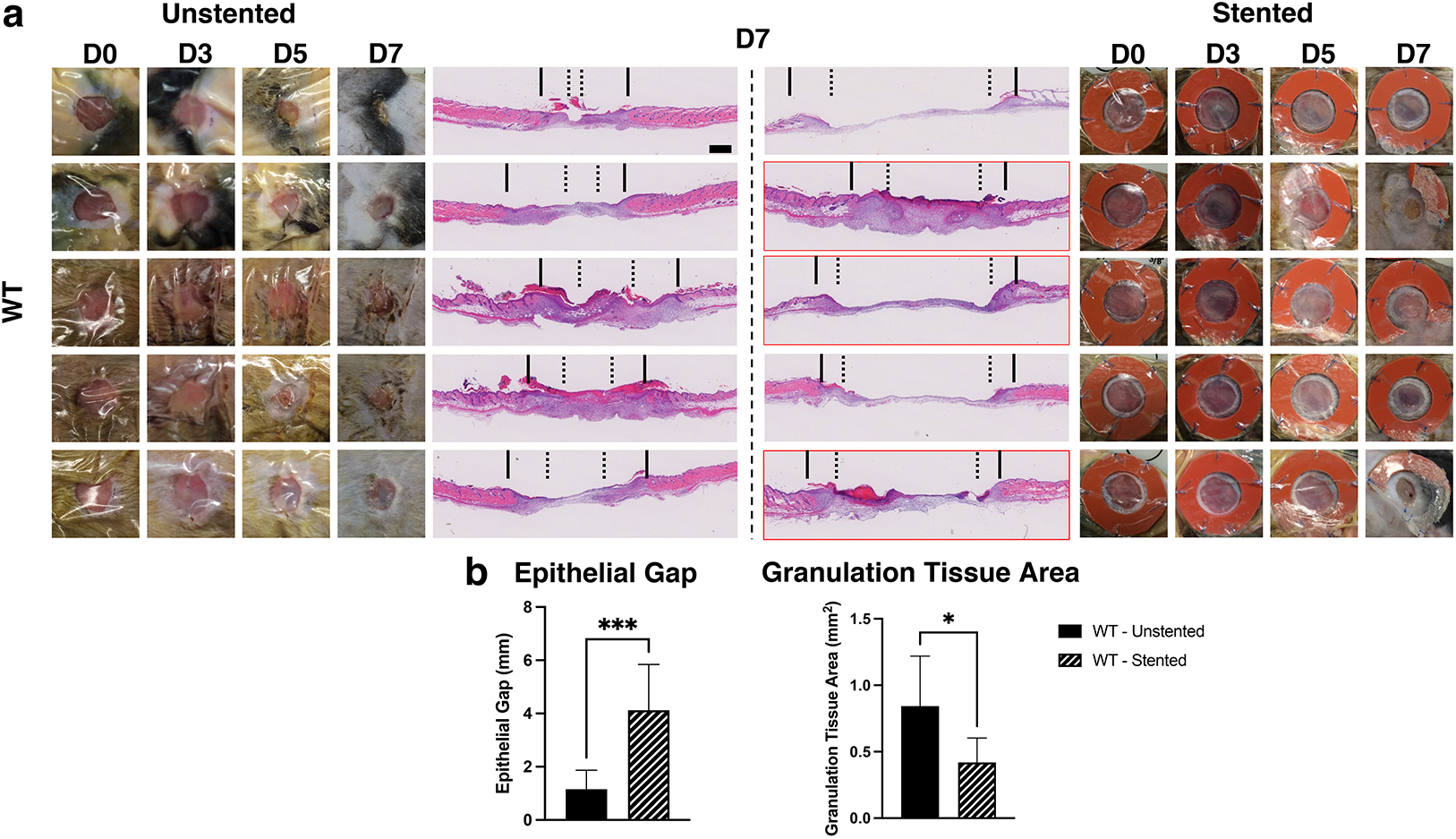
Healing progression of unstented vs stented wounds in WT mice at 7 days post-wounding. **a**. Gross images taken at time of wounding and indicated intervals post-wounding, with representative H&E staining of post-wounding day 7 wounds. Solid vertical line represents original wound margin, dotted line represents extent of re-epithelialization. Red boxes around H&E images represent cases in which stent integrity was compromised prior to day 7. **b**. quantification of epithelial gap and granulation tissue area at day 7 post-wounding. Only wounds in which stents were intact at post-wounding day 7 were included in quantified analysis. Scale bar = 50μm. n=5-10 wounds per treatment group. p-values: *<0.05, **<0.01, ***<0.001, ****<0.0001

### Wounds in WT and IL-10^-/-^ mice heal at the same rate when stented

Next, we compared unstented wound closure of WT and IL-10^-/-^ mice, which we then related to stented wounds of each mouse strain **(Figure2)**. When no stent was applied, but a moist wound environment was provided via a semi occlusive dressing, there was no statistical difference in the epithelial gap at 7 days post-wounding (WT 1.16±0.71 mm vs IL10^-/-^ 1.69±0.93 mm, p=ns) **(Figure 2b)**. Even when stented, no significance was found in epithelial gap (WT 4.13±1.72 mm vs IL10^-/-^ 4.63±1.80 mm, p=ns). When unstented, IL-10^-/-^ mice had greater granulation tissue area compared to WT mice (WT 0.84±0.38 mm^2^ vs IL-10^-/-^ 1.44±0.55 mm^2^, p<0.01). Stenting resulted in a significant reduction in granulation tissue in wounds of IL-10^-/-^ mice (unstented 1.44±0.55 mm^2^ vs stented 0.33±0.15 mm^2^, p<0.001), which was also seen to a lesser degree in WT mice (unstented 0.84±0.38 mm^2^ vs stented 0.42±0.18 mm^2^, p<0.05).

**Figure 2:**
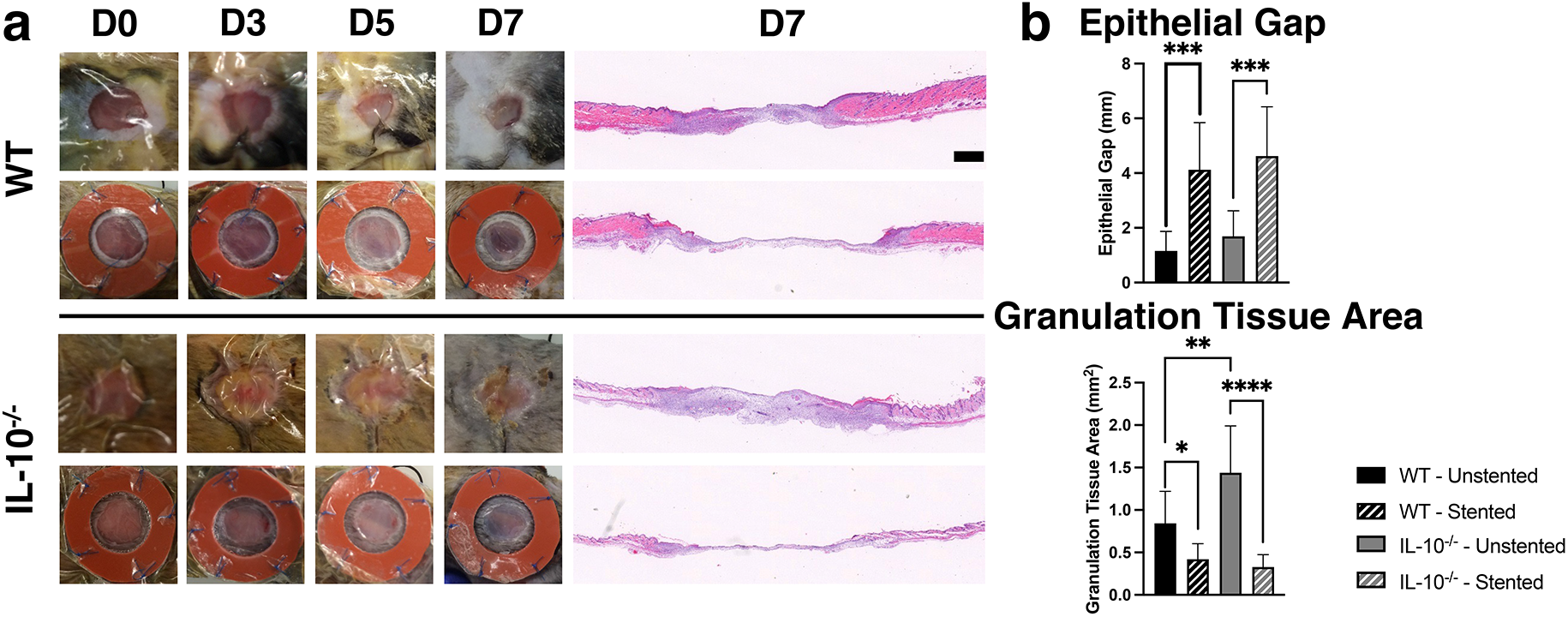
Healing progression of unstented and stented wounds in WT vs IL-10^-/-^ mice at 7 days post-wounding. **a**. Gross images taken at time of wounding and indicated intervals post-wounding, with representative H&E staining of post-wounding day 7 wounds. **b**. Quantification of epithelial gap and granulation tissue area at day 7 post-wounding. Only wounds in which stents were intact at post-wounding day 7 were included in quantified analysis. Scale bar = 50μm. n=5-10 wounds per treatment group. p-values: *<0.05, **<0.01, ***<0.001, ****<0.0001

Additionally, we determined the effect of IL-10 knockout and stenting on myofibroblast activation at day 7 **(Figure 3)**, which tends to follow the course of re-epithelialization. Qualitative examination of total wound α-SMA staining demonstrated that unstented wounds of WT and IL-10^-/-^ both tend to have a focal abundance of α-SMA^+^ myofibroblast presence at the wound edges, which support their assigned role in facilitating wound contraction. The granulation tissue in unstented wounds in IL-10^-/-^ mice appeared to have more α-SMA-positive staining in the middle of the wounds. In comparison, stented wounds in both strains of mice showed a less robust α-SMA-positive myofibroblast abundance at the wound edges at day 7 **(Supplemental Figure 1)**. We quantified the proportion of α-SMA^+^ cells per high power field (HPF) in wounds **(Figure 3b).** No statistically significant difference was noted between genotypes in both the unstented (WT 15.3±6.3% vs IL10^-/-^ 16.4±7.1%, p=ns) and stented (WT 20.1±9.1% vs IL10^-/-^ 16.9±3.7%, p=ns) groups. Similarly, stenting did not significantly alter the proportion of α-SMA^+^ cells within genotypes.

**Figure 3:**
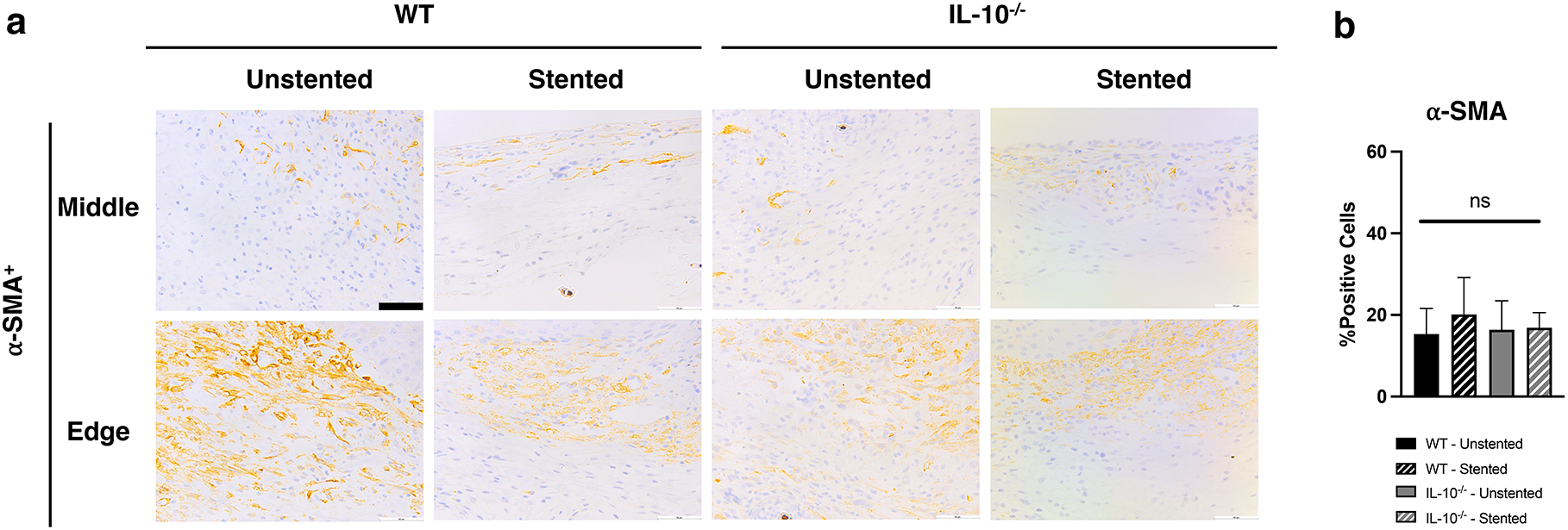
⍰-SMA immunohistochemical staining of unstented and stented wounds in WT vs IL-10^-/-^ mice at 7 days post-wounding. **a**. ⍰-SMA staining representative images from middle and edge of wounds at day 7 post-wounding. **b**. Quantification of % of ⍰-SMA positive cells at day 7 post-wounding. Only wounds in which stents were intact at post-wounding day 7 were included in quantified analysis. Scale bar = 50μm. n=5-10 wounds per treatment group. p-values: *<0.05, **<0.01, ***<0.001, ****<0.0001

### Stenting alters the inflammatory infiltration in the healing wound

We next determined the effect of IL-10 knockout and stenting on inflammatory cell infiltration into wounds at day 7. We determined the leukocyte infiltration in each experimental condition using an anti-CD45 antibody **(Figure 4a–b)**. While there was no statistical significance when stents were applied within both WT (unstented 6.0±2.7% vs stented 10.3±3.5%, p=ns) and IL10^-/-^ (unstented 28.9±14.1% vs stented 22.4±1.5%, p=ns), there was a significantly higher percentage of CD45^+^ cells per HPF in unstented IL10^-/-^ mice compared to unstented WT mice (WT 6.0±2.7% vs IL10^-/-^ 28.9±14.1%, p<0.01).

**Figure 4:**
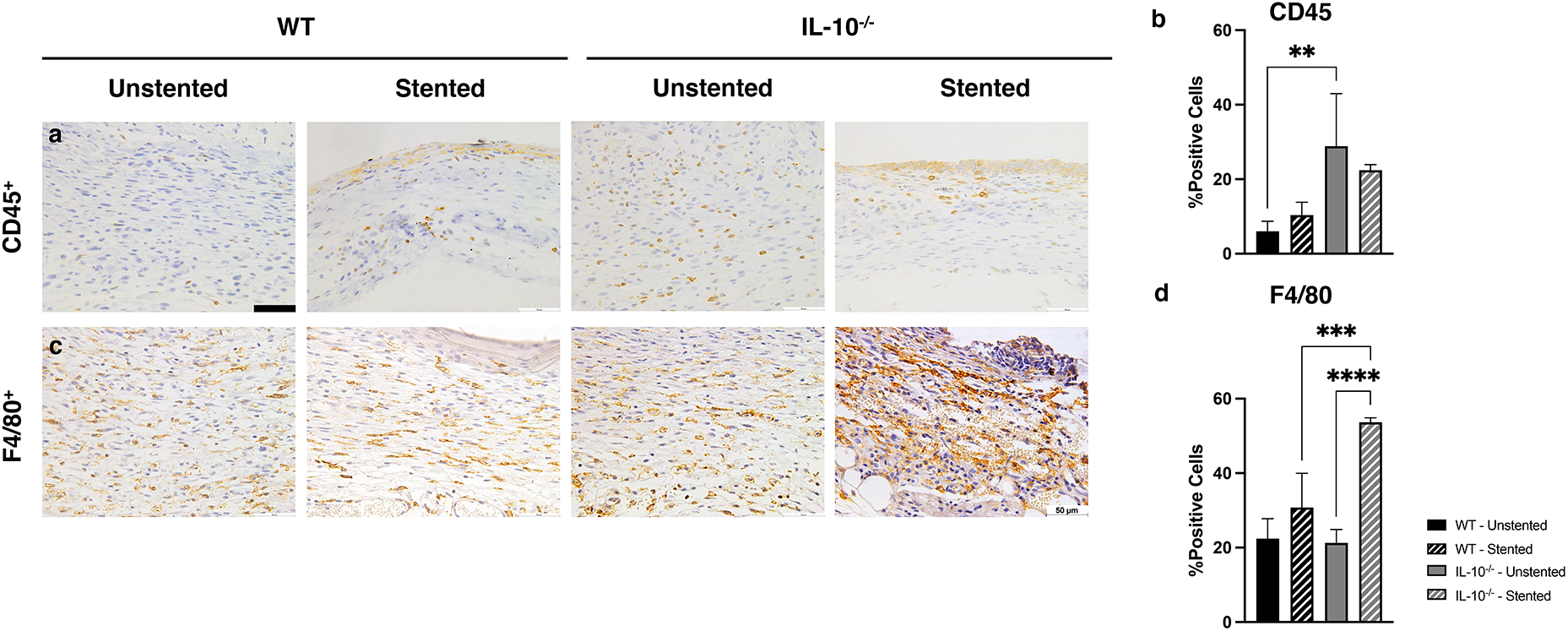
CD45 and F4/80 immunohistochemical staining of unstented and stented wounds in WT vs IL-10^-/-^ mice at 7 days post-wounding. **a**. CD45 staining representative images wounds at day 7 post-wounding. **b**. Quantification of % of CD45 positive cells at day 7 post-wounding. **c**. F4/80 staining representative images wounds at day 7 post-wounding. **d**. Quantification of % of F4/80 positive cells at day 7 post-wounding. Only wounds in which stents were intact at post-wounding day 7 were included in quantified analysis. Scale bar = 50μm. n=5-10 wounds per treatment group. p-values: *<0.05, **<0.01, ***<0.001, ****<0.0001

Quantification of differences in the level of macrophage infiltration was assessed as the percentage of F4/80^+^ cells per HPF **(Figure 4c–d)**. While there was no statistical difference in between genotypes in the non-stented group (WT 22.4±5.4% vs IL10^-/-^ 21.3±3.6%, p=ns), when a stent was applied to the wound, the IL-10^-/-^ mice had a significantly higher percentage of macrophages per HPF (WT 30.8±9.2% vs IL10^-/-^ 53.7±1.1%, p<0.001). Moreover, within the IL10^-/-^ group, stented wounds had a significantly higher level of macrophage infiltration than non-stented wounds (stented 54±10% vs unstented 21±4%, p<0.0001).

### Loss of IL-10 expression results in greater dermal fibrosis

To confirm the effect of loss of IL-10 expression on fibrosis and scarring outcome, we assessed wounds at day 30 post-wounding for dermal architecture **(Figure 5)**. H&E staining of the scars **(Figure 5a&c)** showed an increased density of collagen bundles in the dermis of both unstented and stented IL-10^-/-^ mice scars. While there was no difference in scar length or epithelial thickness across the four groups at the day 30 time point, the stented IL10^-/-^ group had significantly greater scar area than the unstented IL10^-/-^ (unstented 177,220.2± 12,608.1 μm^2^ vs stented 241,660.9±15,147.3 μm^2^, p<0.01), with a similar, yet non-significant, trend observed in the WT cohort (unstented 131,716.4± 28,666.1 μm^2^ vs stented 173,923.7± 61,198.2 μm^2^, p=ns). Moreover, there was a significantly higher measure of epithelial topology in the IL10^-/-^ stented cohort than in the WT stented (WT 215.9± 9.9 μm vs IL10^-/-^ 249.8± 54.3 μm, p<0.05) and IL10^-/-^ unstented groups (unstented 208.2± 4.0 μm vs stented 249.8± 54.3 μm, p<0.01).

**Figure 5:**
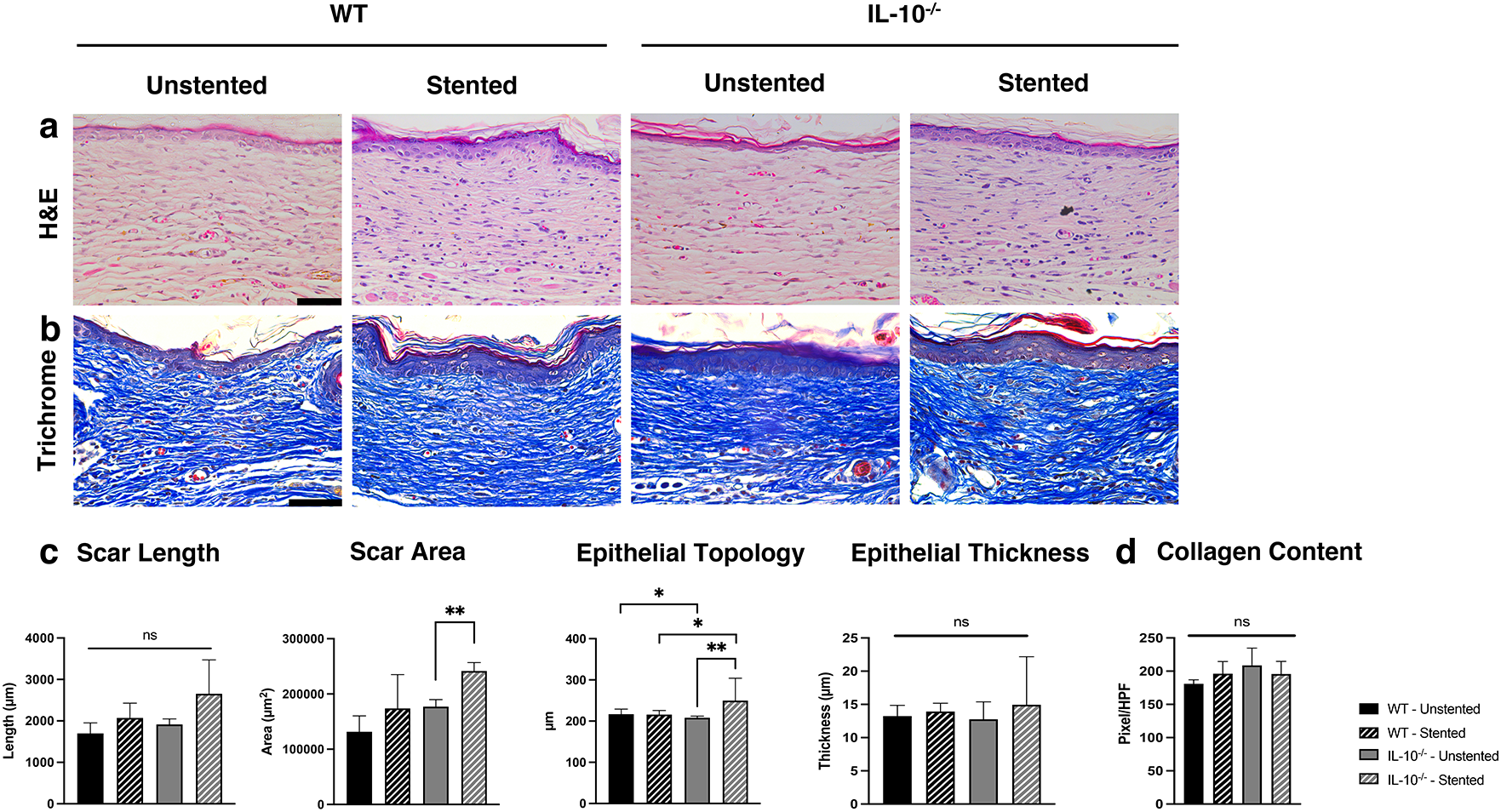
H&E and Trichrome staining of unstented and stented wounds in WT vs IL-10^-/-^ mice at 30 days post-wounding. **a**. Representative images of H&E staining of wounds at day 30 post-wounding. **b**. Quantification of scar measures day 30 post-wounding including scar length and area, epithelial topology and thickness. **c**. Representative images of trichrome staining of wounds at day 30 post-wounding. **d**. Quantification of collagen content at day 30 post-wounding. Only wounds in which stents were intact at post-wounding day 7 were included in quantified analysis. Scale bar = 50μm. n=3-5 wounds per treatment group. p-values: *<0.05, **<0.01, ***<0.001, ****<0.0001

Fibrosis in scars at day 30 was quantified by examining collagen content using trichrome staining (**Figure 5b&d**). We noted dense collagen bundles in dermis of both stented and unstented wounds of the IL-10^-/-^ mice as compared to their WT counterparts. However, upon quantifying pixel density of the blue (collagen positive) staining, there was only a trend indicating a higher collage content in the IL-10^-/-^ mice, with no statistical difference within unstented (WT 180.9±6.1 vs IL-10^-/-^ 208.6±26.2, p=ns) or stented groups (WT 196.2±18.3 vs IL-10^-/-^ 195.8±18.9, p=ns).

## Discussion

Our data indicate that endogenous IL-10 expression does not alter closure of full thickness excisional wounds when controlling for wound hydration and excessive contraction. This suggests that the role of IL-10 in wound healing may be highly dependent on the tissue environment. Factors that might be expected to differ in stented versus unstented mice that could affect the impact of IL-10 include mechanotransduction pathways, cellular metabolism, tissue hydration levels, and others.[26–28] These findings do not invalidate previous work on IL-10 in wound healing, but rather emphasize that the exact models and tissue conditions are important variables. Nonetheless, these data suggest that previous reports of increased rates of re-epithelialization in IL-10^-/-^ mice should be revisited in light of recent advances in wound healing models.[22] We also find that the loss of IL-10 leads to increased inflammatory cell infiltration and scarring. Stenting alters the inflammatory infiltrate of the wound with IL-10^-/-^ mice demonstrating greater inflammatory cell infiltration than WT. This result is consistent with the known role of IL-10 as a regulator of innate and adaptive immune responses suggesting that IL-10 contributes to regulation of inflammation without compromising the healing responses.

Given the role of IL-10 on several local and systemic functions that may modulate cutaneous wound healing, the total loss of IL-10 in this model may affect notable pathways involved in physiologic wound healing. Perhaps of greater impact on wound healing is the effect of IL-10 on macrophage function and polarization. IL-10 is one of many signals driving polarization of macrophages to the alternatively activated, or M2, subset. This macrophage subset is regarded as having anti-inflammatory and regenerative functions, in part mediated by further production of IL-10. Additionally, IL-10 production by macrophages has been implicated in the resolution of the acute wound healing response, resulting in clearance of myofibroblasts from the wound.[29] Since the macrophages in this study lack a major functional property provided by IL-10 secretion, we did not particularly study the M2 subset as the physiologic significance of these macrophages on wound healing and myofibroblast clearance in the setting of no IL-10 production by these cells may not lead to any definitive conclusions [22].

Stenting full thickness wounds for 7 days resulted in dramatic alterations in granulation tissue deposition and prolonged time of epithelial gap closure in both WT and IL-10^-/-^ mice, indicating that the use of stents prevents immediate contraction of dermal wounds in mice. The effect of stenting becomes even more evident when a wound’s stent is compromised, typically by a mouse chewing the stent off, wherein the wound morphology reflects that of unstented wounds perhaps owing to contraction. This is in line with the well-established, significant role that contracture has on wound healing. The method of applying stents to murine wounds to prevent immediate contraction was first introduced by Galiano and colleagues in 2004 as a means to replicate human wound healing.[20] By preventing immediate contraction, this method isolates tissue granulation and re-epithelialization as the means by which these murine wounds heal, which closely replicates the human healing phenotype. Though both the stented and unstented wounds of IL-10^-/-^ mice resulted in greater degrees of scarring compared to their WT controls, the early progression of wound healing was dramatically altered by stenting.

Our work does include several limitations. In our experience, it is common for mice to chew off stents, thus exacerbating the effort and cost of performing these experiments. Even when the stents remain intact for the first 7 days, we presume that every mouse does chew or manipulate their stents in some way. This repeated minor trauma may result in mechanical strain which furthers inflammatory infiltration into and surrounding the wound.[30, 31]. This kind of mechanical tension has also been shown to modulate would healing by stimulating vascular remodeling and cell proliferation in vivo and to lead to dermal fibroblast differentiation in-vitro.[32, 33] Chewing of the stent may be reduced by singly-housing mice following wounding, however this can again add to the cost of experiments and may be precluded by regulations set by local Institutional Animal Care and Use Committees (IACUC). In the same vein, suturing the stent to the skin itself adds potential for foreign body reactions as well as a source of potential bacterial contamination in the skin surrounding the wound. The foreign body reaction is a macrophage driven response defined as cell infiltration, inflammation, and extracellular matrix deposition meant to isolate foreign material from the rest of the body.[34] While this may lead to increased inflammation, it can be seen as comparable to a surgeon closing an incision with suture. Furthermore, we find that stenting WT wounds does not significantly alter macrophage quantity in the wound, while stenting IL-10^-/-^ wounds markedly increases macrophage infiltration. This may be due to the lack of inhibitory stimulus to the innate immune response to the foreign body of suture material or mechanical strain from stent manipulation, which further highlights the importance of the genetic context of animals being studied. Furthermore, our practice of removing stents at day 7 is driven by compliance with standards set by the institutional veterinarians within our IACUC, which may allow wounds to revert to unstented phenotype. However, recent trends in murine wound healing research have left stents on until day of harvest.[35] This practice may influence the final scarring outcome, indicating the value of leaving stent on until day of harvest, while also demonstrating that this does not compromise the health of the wounded animal.

## Conclusion

Our work not only adds to the important literature highlighting the importance of using a stented wound rodent model, but also provides insight to the endogenous role of IL-10 in various contexts of wound healing conditions such as maintenance of a moist wound environment and prevention of contraction via stenting. While previous work demonstrated faster wound healing in unstented wounds lacking IL-10, we demonstrate here that stenting and maintaining a moist wound environment leads to no appreciable difference in wound healing rates in IL-10^-/-^ mice. However, the loss of IL-10 leads to increased inflammatory cell infiltration and scarring. While providing a procedure through which human wound healing can be more accurately modeled, we also show that stenting can significantly alter the inflammatory profile of wounds depending on the genetic background of the model animal utilized. These new findings signify the therapeutic potential for IL-10 overexpression in regulating inflammation and tissue fibrosis without compromising the rate of wound healing.

## Supplementary Materials

Supplemental information may be found at the following link.

## Author Contributions

SB and SGK designed the study; SB, LY, MR, and TL designed and performed experiments; WDS, BP, TP, FF, and OOO analyzed data; WDS, BP, OOO, and FF made the figures; WDS, SB, BP, PB, SGK, and OOO drafted/revised the paper; SB, PLB and SGK provided scientific comments for the paper; and all authors discussed the results and commented on the manuscript.

## Acknowledgements

The authors acknowledge the editorial support of Monica Fahrenholtz, PhD from the Office of Surgical Research Administration at Texas Children’s Hospital.

## Declaration of Competing Interest

The authors declare that they have no competing interests.

## Funding Sources

We acknowledge support for this research by the John S. Dunn Foundation Collaborative Research Award Program administered by the Gulf Coast Consortia, 3M Award given by the Wound Healing Foundation, Clayton Seed Grant by the Department of Pediatric Surgery, Texas Children’s Hospital, and NIGMS-R01GM111808.

## Meeting Presentation

This work was presented at the 17^th^ Annual Academic Surgical Congress in February, 2022 in Orlando, FL.

## Notes

The authors declare no competing financial interests.

### Competing Interest Statement

The authors have declared no competing interest.

