## Supplementary figures and images for "Endogenous IL-10 Contributes to Wound Healing and Regulates Tissue Repair"

### Supplemental Figure 1

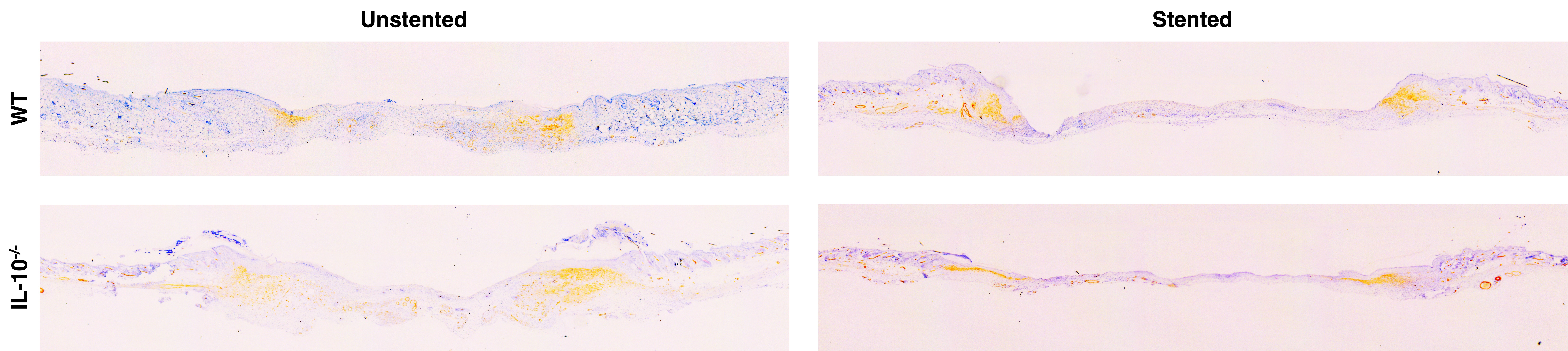
